# Cholesterol Restriction Primes RIG-I Antiviral Responses Through a Noncanonical Type I IFN Pathway

**DOI:** 10.1101/2024.05.19.594902

**Authors:** Tasuku Nishimura, Takahisa Kouwaki, Ken Takashima, Akie Ochi, Yohana S Mtali, Hiroyuki Oshiumi

## Abstract

Cholesterol metabolism is associated with innate immune responses; however, the mechanisms underlying this have not been fully elucidated. Here, we performed a chemical screening to isolate small molecules affecting the activity of RIG-I, a cytoplasmic viral RNA sensor, and found that statins, which inhibit cholesterol synthesis, dramatically enhanced RIG-I-dependent antiviral responses in specific cell types. The restriction of cholesterol synthesis induced the expression of noncanonical type I interferons (IFNs), such as IFN-ω, in an SREBP1 transcription factor-dependent manner. This noncanonical type I IFN expression pathway subsequently enhanced RIG-I-mediated signaling following viral infection. Administration of statins in mice augmented RIG-I-dependent cytokine expression in the lungs. Conversely, a mouse obesity model exhibited reduced RIG-I response in the lungs compared to wild-type mice. Single-cell transcriptome analyses revealed a subset of alveolar macrophages that increased the RIG-I expression in response to inhibited cholesterol synthesis in vivo. This study revealed the noncanonical type I IFN pathway linking cholesterol metabolism and RIG-I signaling. Targeting this pathway could offer valuable insights for developing novel treatment approaches to address future viral pandemics.

## Introduction

The antiviral innate immune response is the first line of defense against viral infections. Recent studies have uncovered the integration of metabolism with immune responses, termed immunometabolism ^1^, and a growing body of evidence has demonstrated that cholesterol metabolism is linked to innate immune responses ^2, 3, 4^. Following the discovery of immunometabolism, recent cohort studies have revealed that metabolic syndromes increase the risk of severity and death in viral infectious diseases ^5^. However, the mechanisms underlying this have not been fully elucidated.

Innate immune responses, including the expression of type I interferon (IFN), are critical for early-phase protection against viral infection ^6, 7, 8, 9^. Viral nucleic acids are recognized by pattern recognition receptors (PRRs), such as RIG-I-like receptors (RLRs), Toll-like receptors (TLRs), and cGAS ^10, 11, 12^. TLRs, including TLR3, 7, and 8, play crucial roles in detecting viral RNAs in endosomes and lysosomes^12^. Conversely, cytoplasmic viral DNAs are targeted by cGAS, which utilizes the STING adaptor molecule to induce type I IFN expression ^11^, while cytoplasmic viral double-stranded RNAs (dsRNAs) are specifically recognized by RLRs, including RIG-I and MDA5 ^10^. RIG-I activation is regulated by several ubiquitin ligases, such as Riplet and TRIM25 ^13,14^. This activation triggers the expression of type I IFNs through the MAVS adaptor molecule, which in turn induces the expression of interferon stimulatory genes (ISGs), including RIG-I itself ^15^. RIG-I senses relatively shorter dsRNAs in the cytoplasm, whereas MDA5 preferentially recognizes longer dsRNAs ^16^. Consequently, RIG-I and MDA5 effectively recognize different types of viruses ^17^. For example, the influenza A virus, Sendai virus (SeV), and several other negative-sense single-stranded RNA viruses are detected by RIG-I. In contrast, picornaviruses (e.g., encephalomyocarditis virus (EMCV) and poliovirus) and coronaviruses (e.g., SARS-CoV and SARS-Cov-2) are primarily recognized by MDA5 ^18, 19, 20^.

Secreted type I IFNs bind to the type I IFN receptors on the cell surface, composed of IFNAR1 and IFNAR2 subunits, which trigger the expression of ISGs and induce antiviral activities ^21^. Several type I IFN genes in the human genome are clustered on chromosome 9, whereas in mice, this cluster is located on chromosome 4. Among these, IFN-α and –β are well-characterized type I IFNs, with more than 10 IFN-α genes encoded in the human genome ^22, 23^. Specifically, IFN-α1 and –α2 expression are highly induced in human peripheral blood mononuclear cells (PBMCs) following infection with several viruses ^24^, and various types of viruses induce IFN-β expression in macrophages as well as epithelial cells and fibroblasts ^25^. In contrast to these canonical type I IFNs, there are several noncanonical type I IFNs, including IFN-ε, –κ, and –ω, as well as minor IFN-α subsets (including IFN-α16). Previous studies have reported that patients with life-threatening COVID-19 possessed neutralizing autoantibodies against IFN-ω and IFN-α ^9^, suggesting the crucial roles of the noncanonical type I IFN. Because noncanonical type I IFNs also stimulate the type I IFN receptor ^21^, their distinct roles remain unclear.

The correlation between metabolic diseases and immune responses to viral infections is noteworthy ^26^, and recent studies have revealed that obesity increases the risk of COVID-19 severity and death ^27, 28^. Moreover, retrospective cohort studies have shown that the use of statins, which inhibit cholesterol biosynthesis, reduces the risk of severity in patients with COVID-19 ^29^. Increasing evidence has begun to clarify the molecular mechanisms that link cholesterol metabolism to antiviral innate immunity. Type I IFN signaling itself regulates cholesterol biosynthesis, and disturbances in cholesterol biosynthesis are associated with an enhanced type I IFN response, in a STING and SREBP2-dependent manner ^2, 3^. The addition of a 20-carbon geranylgeranyl lipid to MAVS by geranylgeranyl transferase type I significantly attenuates type I IFN expression following viral infection, and this geranylgeranylation is associated with cholesterol metabolism ^30, 31^. However, the mechanism through which cholesterol restriction boosts the antiviral response has not been fully elucidated, and the physiological significance of these mechanisms in vivo remains elusive. In this study, we performed a chemical screening to isolate small molecules that enhance RIG-I-dependent type I IFN expression and identified statins. Since statins dramatically enhanced RIG-I-dependent innate immune responses in specific cell types, we aimed to uncover mechanisms linking cholesterol metabolism with the RIG-I-dependent antiviral responses and assess their physiological significance in vivo using mouse models.

## Results

### Statins dramatically enhance RIG-I-dependent antiviral responses

To identify small molecules that modulate RIG-I-mediated type I IFN expression signaling, we developed a chemical screening assay system in which the activation of RIG-I was induced by ectopic expression of the Riplet protein, a coreceptor of RIG-I ^32^. RIG-I/Riplet signaling leads to enhanced mCherry reporter protein expression, driven by IFN-β promoter, with AcGFP, driven by EF1α promoter, serving as an internal control. Our initial screening of approximately 1,600 approved compounds identified 22 molecules (Figure 1A). The second screening was conducted using the plasmid carrying luciferase driven by the IFN-β promoter (p125-luc) to eliminate artificial fluorescence, and we identified several statins, including pitavastatin, cerivastatin, and atorvastatin, which are known inhibitors of cholesterol synthesis, and found that those statins enhanced RIG-I signaling. We confirmed that pitavastatin treatment significantly increased the activation of the IFN-β promoter by RIG-I in a dose-dependent manner in HEK293 cells (Figure 1B).

**Figure 1.**
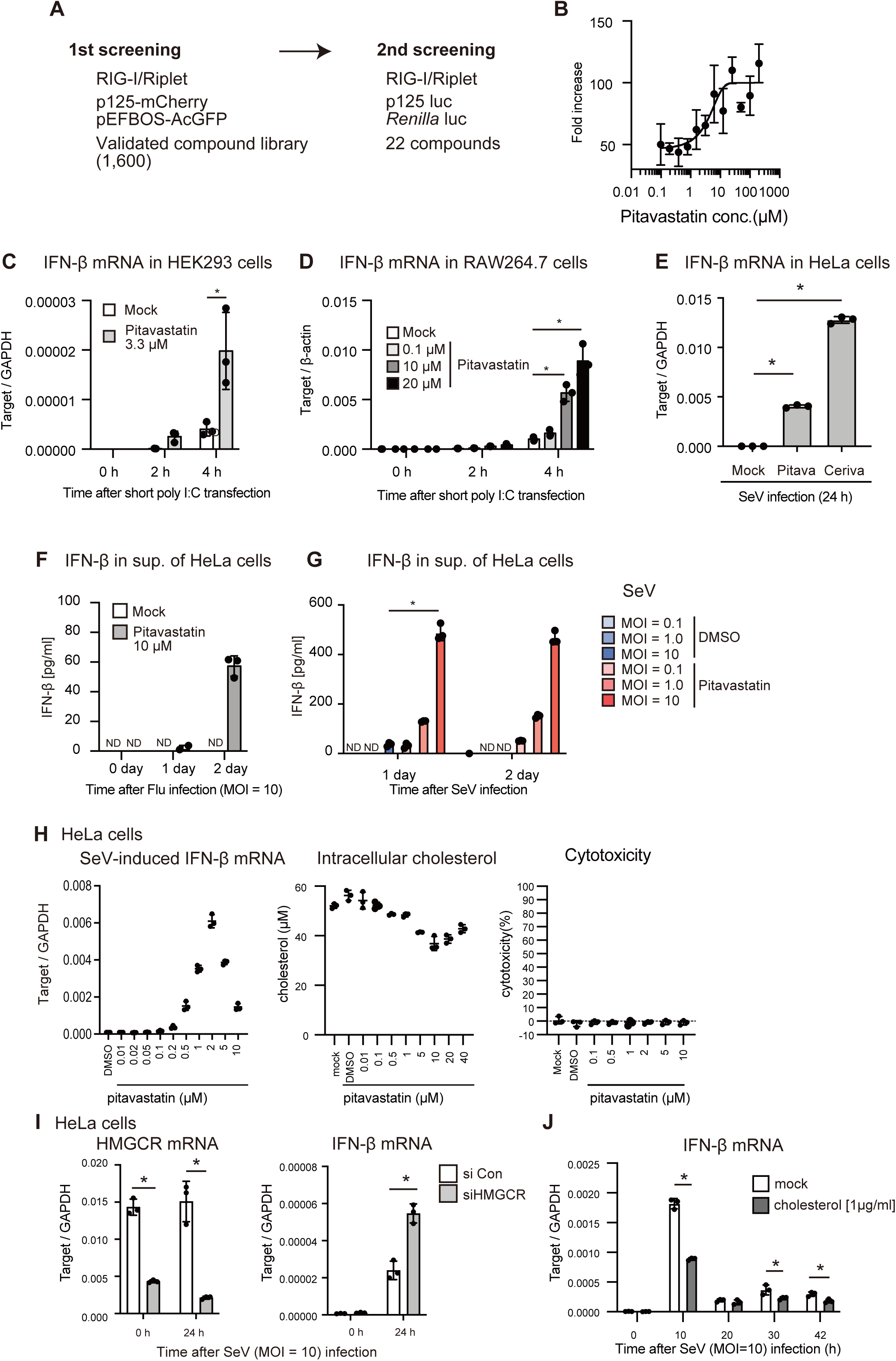
Inverse correlation between cellular cholesterol levels and RIG-I-dependent type I IFN expression levels. A) Schematic representation of chemical screening. B) HEK293 cells were transfected with the p125-luc reporter (IFN-β promoter) and *Renilla* luciferase (internal control) together with RIG-I– and Riplet-expressing vectors. Pitavastatin was added to the cell-culture medium at the indicated concentrations (final concentration) and incubated for 24 h, and the luciferase activity was evaluated. C–F) HEK293, RAW264.7, and HeLa cells were treated with pitavastatin at the indicated concentrations, and subsequently stimulated by short poly I:C transfection. The expression of IFN-β mRNA was determined by RT-qPCR and normalized to that of GAPDH or β-actin as indicated. G) HeLa cells were treated with pitavastatin or DMSO (mock) for 1 day and subsequently infected with the Sendai virus (SeV) for 1 or 2 days. IFN-β protein levels in cell culture sup. were determined by ELISA. H) HeLa cells were treated with pitavastatin at the indicated concentrations for 24 h, and cellular cholesterol levels (center panel) and cytotoxicity (right panel) were determined. Pitavastatin-treated HeLa cells were infected with SeV for 24 h, and IFN-β mRNA levels were determined by RT-qPCR (left panel). I) HeLa cells were transfected with siRNAs targeting HMGCR or control siRNA. After 2 days, the cells were infected with SeV. The mRNA expression of HMGCR and IFN-βwas determined by RT-qPCR and normalized to that of GAPDH. J) Cholesterol was added to HeLa cell culture medium for 24 h, and the cells were infected with SeV. The expression of IFN-β mRNA was determined by RT-qPCR and normalized to that of β-actin.

To further assess the effects of statins on RIG-I signaling, we measured the mRNA expression and protein production of cytokines after RIG-I activation in statin-treated cells. Short poly I:C is a ligand of RIG-I, and treatment of cells with pitavastatin or cerivastatin markedly enhanced the expression of IFN-β, IL-6, and IP10 mRNA in HEK293, RAW264.7, and HeLa cells following short poly I:C transfection or infection with SeV, which is recognized by RIG-I (Figures 1C–E and EV1A–C). Pitavastatin treatment dramatically increased the levels of IFN-β proteins produced by HeLa cells in response to infection with influenza A virus (Flu) or SeV (Figures 1F and G). These data suggested that pitavastatin treatment markedly enhanced RIG-I-mediated cytokine expression in HeLa cells.

Next, we assessed the effects on the other PRRs. The addition of poly I:C to the cell culture medium leads to TLR3 activation ^33^, and the mRNA expression of IFN-β and IFN-L3 was enhanced by pitavastatin treatment (Figures EV1D and E), implying that the effect of pitavastatin is not specific to RIG-I signaling. In contrast, cGAMP, a ligand of STING, failed to increase the expression of IFN-β mRNA in HeLa cells, irrespective of the presence or absence of pitavastatin (Figure EV1F), implying the cGAS/STING-independent mechanism.

Considering that pitavastatin inhibits cholesterol synthesis, we hypothesized that cellular cholesterol levels would influence RIG-I-mediated innate immune responses. Firstly, we compared cellular cholesterol levels with the expression levels of type I IFN at various pitavastatin concentrations to evaluate this hypothesis. The results showed that around 1–5 μM of pitavastatin was sufficient to reduce cellular cholesterol levels in HeLa cells, and this same pitavastatin concentration could enhance type I IFN expression following SeV infection (Figure 1H). Furthermore, we confirmed that these concentrations did not induce cytotoxicity (Figure 1H, right panel). This correlation was also observed in HepG2 and A549 cells (Figures EV1H and I), although the effective concentrations differed between these cell lines. A similar correlation was observed when cerivastatin was used instead of pitavastatin (Figure EV1J). To further explore the correlation between cellular cholesterol levels and type I IFN expression, we performed knockdown experiments using siRNA targeting HMGCR, a target of statins. The siRNA for HMGCR markedly reduced HMGCR mRNA and enhanced RIG-I-dependent IFN-β mRNA expression in HeLa and HepG2 cells (Figures 1I, EV1K, and L). Conversely, the addition of cholesterol to the cell culture medium reduced SeV-induced IFN-β expression (Figure 1J). These data indicate that cellular cholesterol levels are inversely correlated with type I IFN expression levels following RIG-I stimulation.

### Noncanonical type I IFN expression are induced by statin treatment

To investigate the mechanism underlying these above findings, we performed a microarray analysis of HeLa cells before and after SeV infection with or without pitavastatin treatment. We confirmed that pitavastatin treatment enhanced the expression of genes essential for lipid synthesis, including cholesterol synthesis (Figure 2A). Following this, we focused on the expression of genes involved in the innate immune response. Interestingly, several noncanonical type I IFN genes, including those for IFN-W1 (IFN-ω), IFN-A4 (IFN-α4), IFN-A5, IFN-A16, IFN-A17, IFN-L3, and IFN-E, as well as IRF transcription factors, and RLRs, were moderately upregulated by pitavastatin pretreatment in the absence of SeV infection (Figures 2B and C). In contrast, IFN-B1, a canonical type I IFN, was markedly upregulated only after SeV infection (Figure 2B). These data suggest that pitavastatin treatment leads to the expression of noncanonical type I IFNs and RLRs before viral infection.

**Figure 2.**
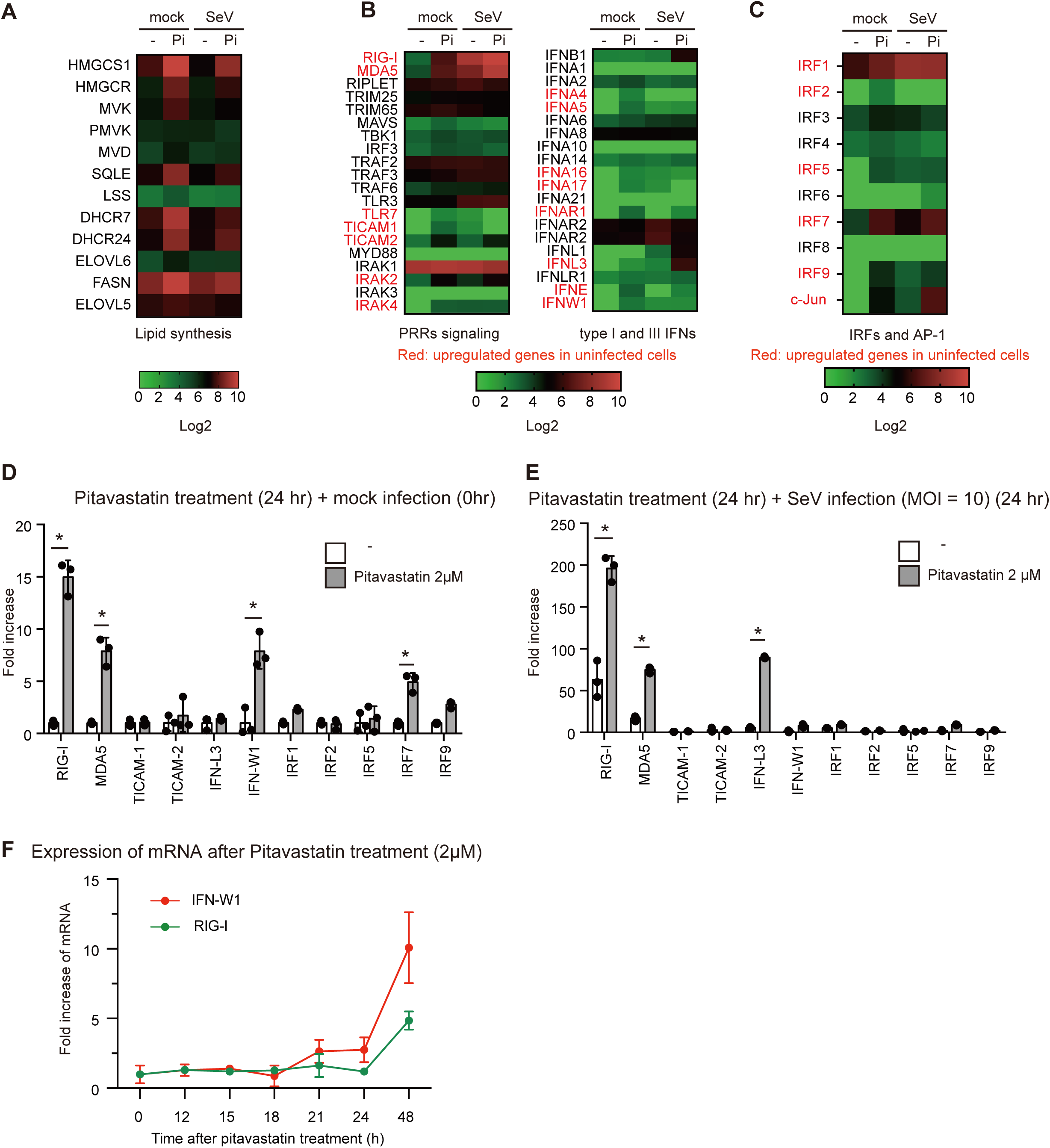
Pitavastatin-induced expression of RIG-I and non-canonical type I IFNs in the absence of cell stimulation. A-C) Heat maps of gene expression in pitavastatin-treated HeLa cells with or without SeV infection. HeLa cells were treated with pitavastatin (Pi) for 24 h, followed by infection with SeV or a mock treatment. Gene expression profiles of each sample were determined by a microarray analysis. D, E) HeLa cells were treated with pitavastatin for 24 hr, followed by infection with SeV (E) or mock treatment (D) for 24 h. The expression of each gene was determined by RT-qPCR and normalized to that of β-actin. The fold increase was calculated by dividing the normalized expression level of a pitavastatin-treated sample by the corresponding value of a non-treated sample. F) HeLa cells were treated with pitavastatin or a solvent (DMSO), and the expression of RIG-I and IFN-W1 at the indicated time points was determined by RT-qPCR and normalized to that of β-actin. The fold increase was calculated by dividing the mRNA levels of pitavastatin-treated cells by those of DMSO-treated cells.

RT-qPCR was conducted to confirm the results of the microarray analysis. The increases in the expression of RIG-I, MDA5, and IFN-W1 mRNA expression following pitavastatin treatment in the absence of SeV infection were statistically significant (Figure 2D). Furthermore, pitavastatin treatment also significantly increased IFN-A16 mRNA expression without SeV infection (Figure EV2A). The increase in IFN-W1 expression following SeV infection was moderate, whereas RIG-I and MDA5 expression were markedly increased following SeV infection (Figure 2E). Pitavastatin treatment also increased IFN-W1 mRNA levels in HepG2 cells but not in A549 and THP-1 cells at the experimental condition (Figure EV2B). Because RIG-I is an interferon-inducible gene, we compared the kinetics of IFN-W1 and RIG-I gene expression after pitavastatin treatment and found that RIG-I expression was delayed compared with IFN-W1 expression (Figure 2F), suggesting that RIG-I expression occurs following IFN-W1 expression.

Based on the above data, we anticipated that the initial expression of noncanonical type I IFNs would induce subsequent RIG-I expression. To evaluate this hypothesis, cells were treated with the IFN-ω recombinant proteins, and RIG-I expression was measured. As expected, stimulation with the IFN-ω protein markedly induced the RIG-I expression (Figure 3A). However, IFN-β mRNA expression was hardly induced by the IFN-ω protein (Figure 3B). Recombinant IFN-ω protein also increased the expression of IRF7, MDA5, and TLR3, but not cGAS mRNAs (Figures 3C-F). Adding 10 pg/mL of recombinant IFN-ω protein was sufficient to enhance the expression of IFN-β and RIG-I mRNA following short polyI:C stimulation, similar to the effect of pitavastatin (Figures 3G and EV3A). If pitavastatin treatment increases RIG-I expression via noncanonical type I IFNs, it is expected that pitavastatin-induced RIG-I expression will be impaired by a defect in the type I IFN receptor. IFNAR2 is an essential component of the type I IFN receptor, and we generated IFNAR2 knockout (KO) cells using the CRISPR-Cas9 system. When cells were treated with pitavastatin for 24 h, IFNAR2 KO abolished pitavastatin-induced RIG-I expression (Figure 3H), suggesting that the type I IFN receptor was involved in pitavastatin-induced RIG-I mRNA expression. However, 48 h treatment with pitavastatin increased RIG-I expression even in IFNAR KO cells (Figures 3I), implying the existence of both type I IFN receptor-dependent and –independent pathways. Notably, pitavastatin-mediated enhancement of IFN-β mRNA expression following Flu infection was markedly reduced by IFNAR2 KO (Figure 3J). These data are consistent with our hypothesis that initially produced noncanonical type I IFNs, including IFN-ω, induce subsequent RIG-I expression via the type I IFN receptor.

**Figure 3.**
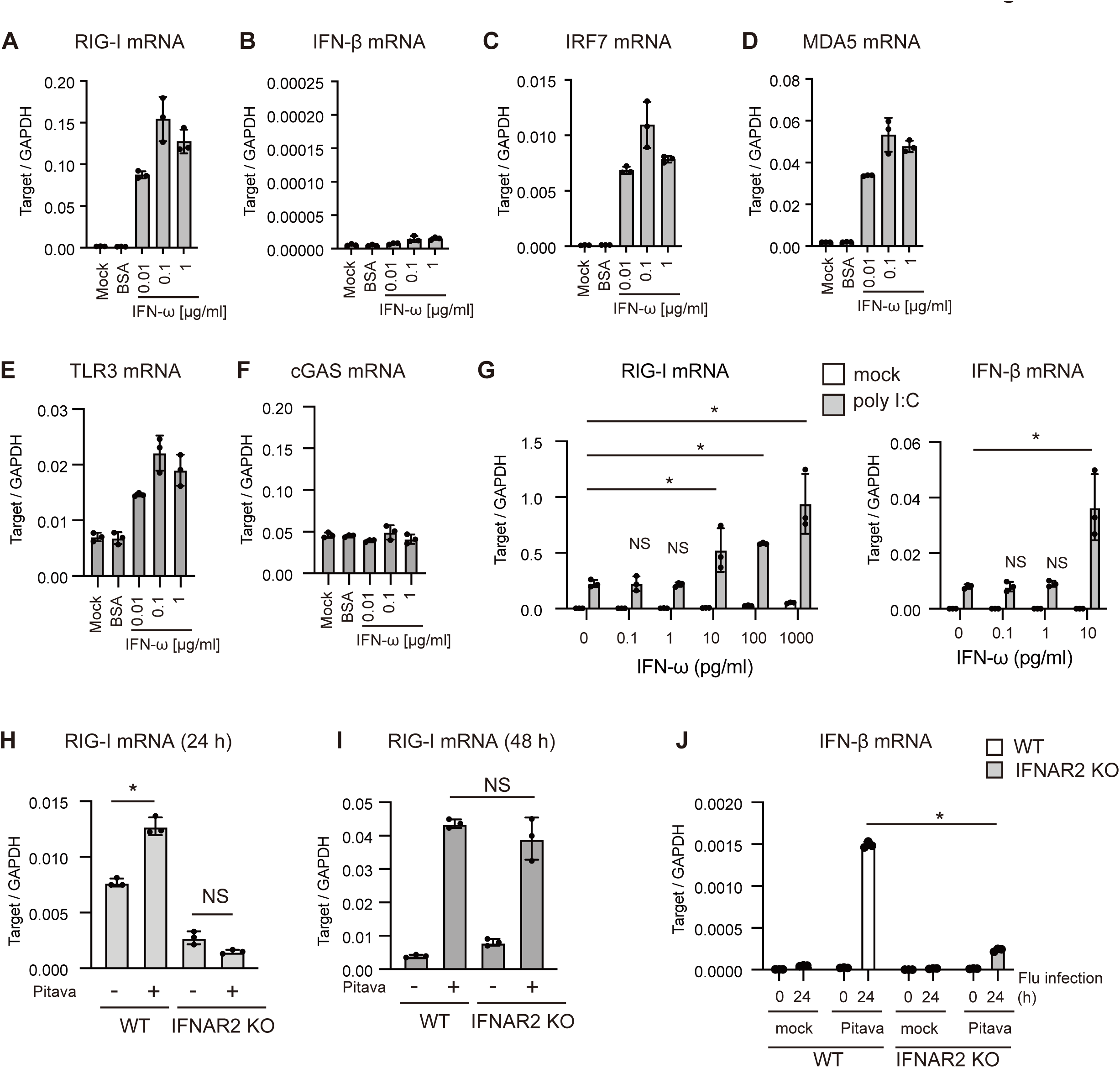
The type I IFN receptor is involved in statin-mediated enhancement of RIG-I signaling. A-F) HeLa cells were treated with recombinant IFN-ω protein at the indicated concentration for 24 h, and expression of each gene was determined by RT-qPCR and normalized to that of GAPDH. G) HeLa cells were treated with recombinant IFN-ω protein at the indicated concentration for 24 hr, and subsequently stimulated via short poly I:C transfection. 4 hr after transfection, IFN-β mRNA expression was determined by RT-qPCR and normalized to that of GAPDH. H, I) Wild type (WT) and IFNAR2 knockout (KO) HeLa cells were treated with or without pitavastatin for 24 (H) or 48 h (I), and the expression of RIG-I was determined by RT-qPCR and normalized to that of GAPDH. J) WT and HeLa cells were treated with pitavastatin (Pitava) or DMSO (mock) for 24 hr, and then infected with Flu. The expression of IFN-β mRNA at indicated time points was determined by RT-qPCR and normalized to that of GAPDH.

### The noncanonical type I IFN expression pathway is dependent on the SREBP1 transcription factor

A decrease in cellular cholesterol levels is known to induce the activation of the transcription factors, SREBP1 and SREBP2 ^34^. Therefore, we investigated the roles of SREBP1 and SREBP2 in this process. First, we measured promoter activity using reporter assay with constitutively active forms of the SREBPs (SREBP1a-N, 1c-N, and 2-N) and RIG-I (RIG-I CARDs). Ectopic expression of RIG-I CARDs markedly activated the IFN-β promoter but not the IFN-ω promoter (Figure 4A). In contrast, ectopic expression of SREBP1a-N, 1c-N, and 2-N markedly activated the IFN-ω promoter (Figure 4A). The IRF7 promoters were also activated by SREBP1a-N and 2-N, but not by RIG-I CARDs (Figure 4A). The RIG-I promoter itself was not activated by the ectopic expression of SREBP1a-N, 1c-N, and 2-N, although the promoter was effectively activated by treatment with IFN-α (Figure 4A). The ectopic expression of MAVS, MyD88, and STING activated the IFN-β promoter but hardly activated the IFN-ω promoter, while SREBP1a-N induced the activation of the IFN-ω promoter (Figure EV4A). These data suggested that the regulation of IFN-ω expression was distinct from that of IFN-β expression.

**Figure 4.**
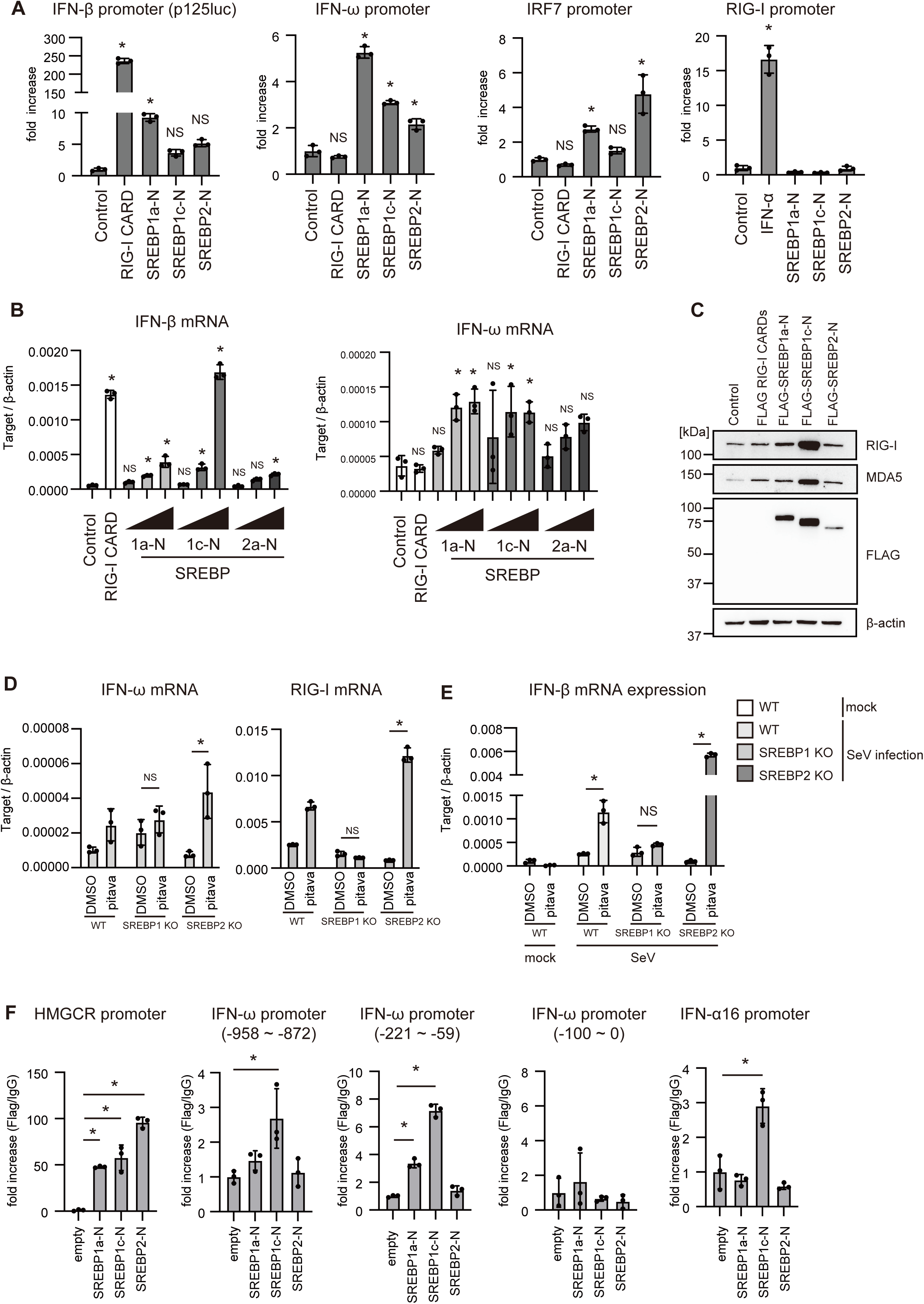
SREBP1 is essential for stain-mediated enhancement of RIG-I signaling. A) HEK293 cells were transfected with either the reporter plasmid for each gene promoter along with the expression vector expressing the indicated gene and an empty vector (control) or stimulated with 100 U of IFN-α. 24 hr after transfection or stimulation, cells were lysed, and the activity of each promoter was determined by measuring luciferase activities. The fold increase was calculated by dividing the luciferase activity of each sample by the average luciferase activity of the control samples. B, C) HEK293 cells were transfected with 200 ng/mL of RIG-I CARD expressing vectors along with 200, 1000, or 2000 ng/mL of SREBP1a-N, 1c-N, or 2a-N expressing vectors in a 24-well plate. The total amount of DNAs was normalized to the same amount by adding the empty vector. 24 h after transfection, the expression of each gene was determined by RT-qPCR and normalized to that of β-actin (B), or cell lysates were prepared and subjected to SDS-PAGE and stained by western blotting with the indicated antibodies (C). D, E,) WT, SREBP1 KO, and SREBP2 KO HeLa cells were treated with pitavastatin (pitava) and solvent (DMSO) for 24 h. Following this, the expression of each gene was determined by RT-qPCR and normalized to that of β-actin (D), or treated cells were infected with SeV for 24 hr, and the expression of IFN-β mRNA was determined by RT-qPCR and normalized to that of β-actin (E). F) FLAG-tagged SREBP1a-N, SREBP-1c-N, and SREBP2-N were transfected into HEK293 cells. A chromatin immunoprecipitation (ChIP) assay was performed using anti-FLAG antibodies. The binding of the protein to the indicated DNA regions was detected by qPCR.

Next, we measured the expression of endogenous IFN-β and IFN-ω genes. Ectopic expression of RIG-I CARDs induced the mRNA expression of IFN-β, along with that of IP-10, IRF7, and RIG-I but not the mRNA expression of IFN-ω (Figures 4B, EV4B-E). In contrast, ectopic expression of SREBP1a-N and SREBP1c-N increased the mRNA expression of IFN-ω and those of RIG-I, and HMGCR (Figures 4B and EV4B– E) as well as RIG-I protein levels (Figures 4C). Moreover, short poly I:C stimulation of HeLa cells failed to increase the mRNA expression of IFN-ω but did increase IFN-β mRNA expression (Figure EV4F), although it could induce IFN-ω expression in THP-1 cells (Figure EV4G). These data indicate that SREBP1 and SREBP2, but not RIG-I, have the potential to activate the promoter of the IFN-ω gene.

To further assess the role of SREBPs on IFN-ω expression, we generated SREBP1 and SREBP2 KO cells using the CRISPR-Cas9 system. Pitavastatin-induced IFN-ω expression was abolished by SREBP1 KO but not by SREBP2 KO (Figure 4D). Induced expression of RIG-I mRNA by pitavastatin treatment was also abrogated by SREBP1 KO (Figure 4D). Moreover, SREBP1 KO abolished the effect of pitavastatin treatment on RIG-I-dependent IFN-β mRNA expression following SeV infection (Figure 4E). These findings indicate that SREBP1 is responsible for the effect of pitavastatin on RIG-I signaling.

To confirm the involvement of SREBP1 in the expression of noncanonical type I IFNs, we performed a chromatin immunoprecipitation (ChIP) assay. The SRE and E-box motifs serve as the binding sites for SREBPs ^35^ and are located in the promoter regions of IFN-ω and IFN-α16 (Figures EV4H and I). The ChIP assay showed that SERBP1a-N and SREBP1c-N could bind to the IFN-ω promoter as well as the IFN-α16 promoter; however, SREBP2-N failed to bind to the IFN-ω and IFN-α16 promoters (Figure 4F). These data support the model that SREBP1 is involved in noncanonical type I IFN expression.

### Effect of cholesterol restriction on antiviral innate immunity in vivo

Next, we assessed whether the restriction of cholesterol synthesis augments RIG-I-dependent type I IFN expression in vivo. Pitavastatin was intraperitoneally administered to mice every three days, followed by the intraperitoneal injection of poly I:C. Since the IFN-W1 gene is not conserved in mice, we examined the expression of all the other type I IFNs, as well as RIG-I. Although we could not detect the significant increase of the expression of RIG-I and type I IFNs before poly I:C injection in the lung, pitavastatin administration augmented the expression of RIG-I, IFN-α1, IFN-α16, and several IFN-α subtypes in the lung following poly I:C injection (Figures 5A and EV5A). The augmented expression of RIG-I and type I IFNs before and after poly I:C stimulation was detected in the heart (Figure 5B). However, this augmentation was not observed in the spleen, liver, muscle, or uterus (Figures 5C and EV5A), suggesting tissue specificity.

**Figure 5.**
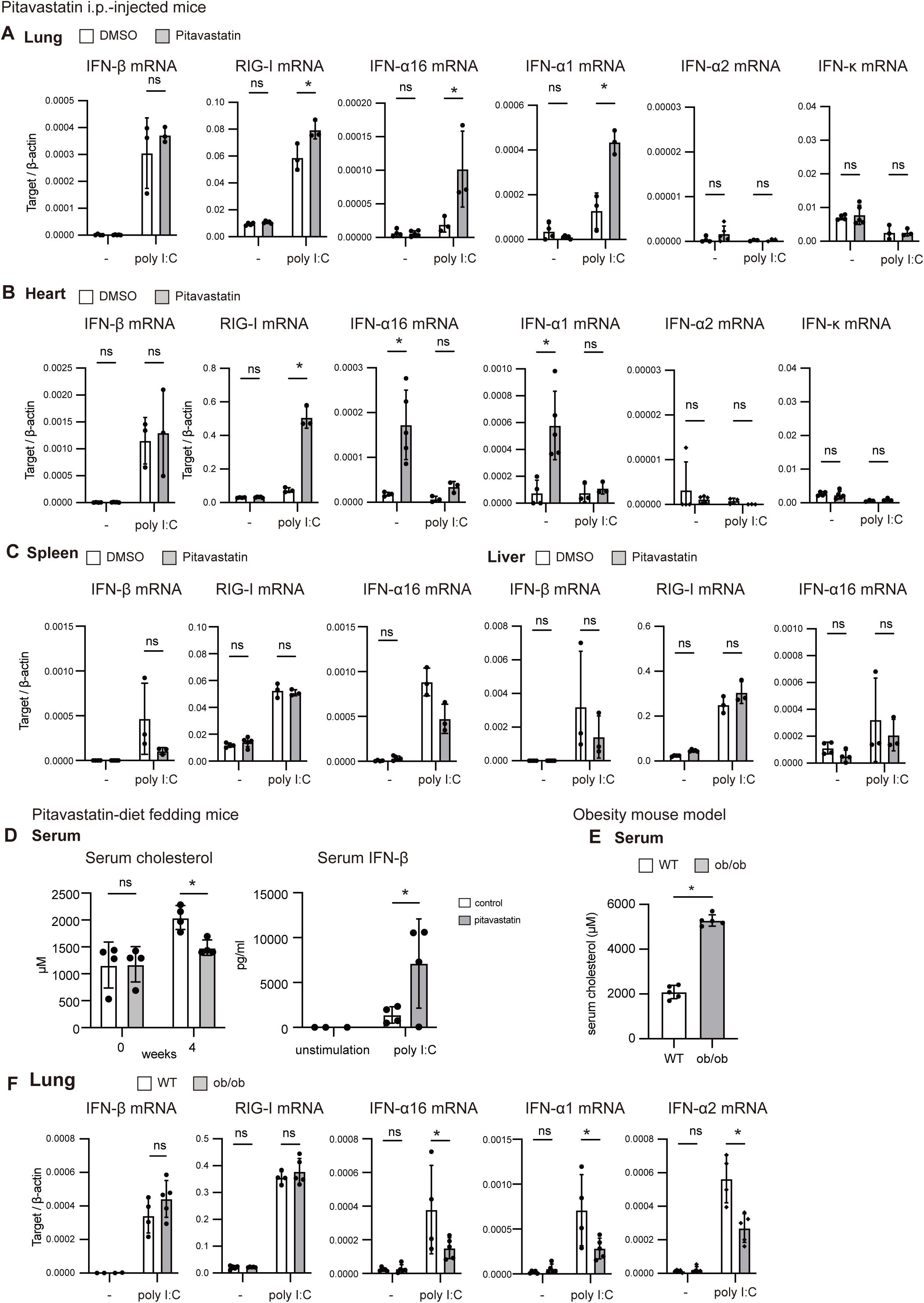
The tissue-specific effects of pitavastatin on RIG-I and type I IFN gene expression. A–C) Pitavastatin or solvent (DMSO) was intraperitoneally injected into mice every day for three days, followed by an intraperitoneal injection of poly I:C or solvent (DMSO). 24 h after poly I:C injection, the lungs (A), heart (B), and spleen (C) were isolated. The total RNA was extracted from each tissue sample. The expression of each gene was determined by RT-qPCR and normalized to that of β-actin. D) Mice were fed a normal or pitavastatin-containing diet for four weeks, following which they received intraperitoneal injection of poly I:C. Sera were collected at each time point, and serum cholesterol and IFN-β protein levels were determined. E, F) Sera from WT and ob/ob mice were collected, and serum cholesterol levels were determined (E). Poly I:C was intraperitoneally injected into WT and ob/ob mice. 6 h after injection, the mRNA expression of RIG-I and type I IFNs mRNA in the lung was determined by RT-qPCR and normalized to that of β-actin.

Next, we used another mouse model, in which mice were fed a pitavastatin-containing diet for 4 weeks, after which they were injected with poly I:C. We confirmed that this treatment reduced blood cholesterol levels (Figure 5D). Notably, the pitavastatin-containing diet markedly augmented serum IFN-β protein levels after poly I:C injection (Figure 5D).

Following this, we used a mouse model of obesity (ob/ob mouse), and we confirmed that serum cholesterol levels were higher in the ob/ob mice, which spontaneously developed obesity ^36^, than those in wild-type mice (Figure 5E). Interestingly, IFN-α1, –α2, and –α16 expression in the lung after poly I:C injection was attenuated in ob/ob mice compared with WT mice (Figure 5F). Collectively, these data suggest that cholesterol levels are inversely correlated with the expression of polyI:C-induced type I IFN and RIG-I in the lungs.

### Identification of CD140a^+^ alveolar macrophages priming the RIG-I pathway in response to cholesterol restriction

In the heart, we detected the increase in the expression of a noncanonical type I IFN, IFN-α16, by pitavastatin treatment without poly I:C stimulation. However, induced expression of noncanonical type I IFNs and RIG-I by pitavastatin treatment alone was not detectable in the lungs of the above animal models. Therefore, we anticipated that only a small subset of lung cells expressed RIG-I in response to pitavastatin administration before poly I:C injection. We expected that single-cell transcriptome analysis of the mouse lung would reveal the RIG-I expression in response to inhibited cholesterol synthesis in specific cell subtypes.

To investigate cell populations that express RIG-I in response to pitavastatin administration, we conducted a single-cell RNA sequence (scRNA-seq) analysis. Single-cell suspensions of whole lungs were prepared from mice injected with DMSO (n =3) or pitavastatin (n = 3) (Figure EV6A). After quality control, a total of 41,259 cells (DMSO group: 18,415 cells and pitavastatin group: 22844 cells) were retained for downstream analysis (see the Materials and Methods section for details). UMAP analysis revealed a diverse array of lung cell populations, identifying numerous distinct cell types, such as alveolar epithelial cells, macrophages, T cells, and endothelial cells (Figure EV6B). Comparative UMAP plots showed the lung cell populations in DMSO– and pitavastatin-administered mice and suggested that the populations of each cell type were largely unaffected by pitavastatin treatment (Figure EV6C). We then focused on cells expressing genes involved in cholesterol metabolism (GO: 0008203) and type I IFN production (GO: 0032606). Accordingly, we found that the cluster identified as alveolar macrophages expressed genes were associated with cholesterol metabolism and type I IFN production (Figure EV6D), prompting a more detailed investigation of the alveolar macrophage cluster.

Single-cell transcriptomics was employed to investigate the heterogeneity of alveolar macrophages within the lung tissue and to elucidate the response to pitavastatin treatment (Figure 6A). Sub-clustering of alveolar macrophages revealed the presence of 11 distinct subpopulations, each characterized by a unique transcriptional profile (Figure 6A-C). This stratification allowed for a detailed examination of gene expression variations within these populations. The violin plots in Figure 6D provide a graphical representation of the expression levels of selected genes implicated in the innate immune response across the different alveolar macrophage subclusters. Notably, RIG-I (DDX58) expression was prominent in clusters 6, 8, and 10, while cGAS was predominantly expressed in cluster 5 (Figure 6D). Interestingly, cluster 8 expressed CD11b (Itgam), and cluster 5, characterized by Mki67, indicating cell division, was predominantly associated with the G2/M phase of the cell cycle, as shown in the UMAP plots (Figure 6D and E). This suggested that cGAS-expressing alveolar macrophages were proliferative.

**Figure 6.**
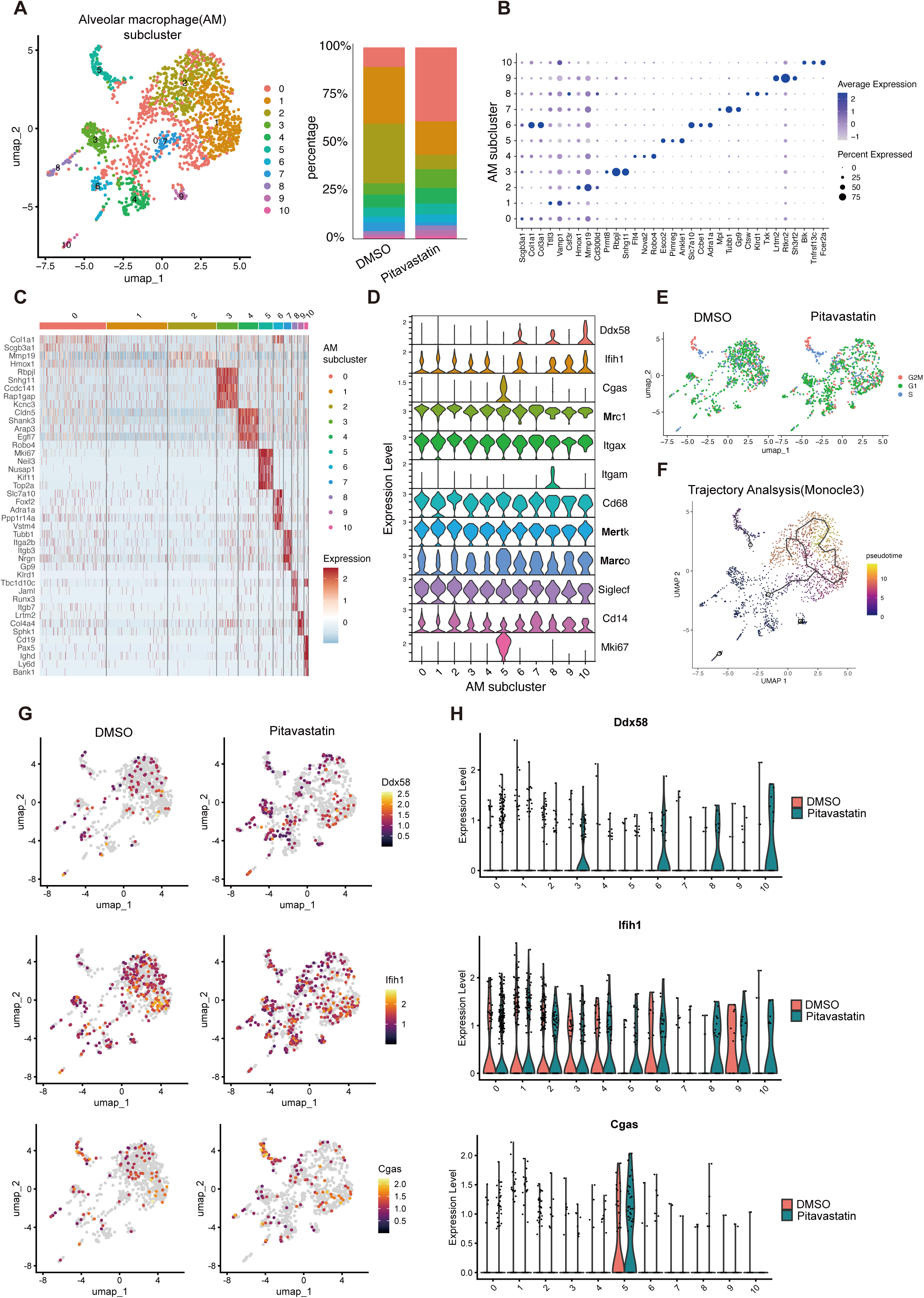
Detailed single-cell RNA Seq analysis of alveolar macrophage subpopulations in the lungs of pitavastatin-treated mice. A) UMAP plots of alveolar macrophage subclusters identified within the lung cell populations. Each dot represents individual alveolar macrophages, with distinct colors indicating the different subclusters. The bar chart on the right shows the proportion of each subcluster in pitavastatin or DMSO-treated mice. B) Dot blot illustrating the expression levels of selected genes in various alveolar macrophage subclusters. The size of each dot corresponds to the percentages of cells expressing the gene within each subcluster, while the color intensity indicates the average expression levels. C) Heatmap of gene expression across alveolar macrophage subclusters, indicating the scaled expression level of a subset of genes. D) Violin plots depicting the expression distribution of selected genes among alveolar macrophage subclusters including both DMSO and pitavastatin-treated groups. The expression level (log-normalized) is shown on the y-axis. E) Comparative UMAP plots showing cell cycle annotation of alveolar macrophages from DMSO– and pitavastatin-treated mice. F) Trajectory analysis (Monocle 3) displaying the pseudo-time of alveolar macrophages, suggesting a developmental or activation trajectory. G) UMAP plots showing the spatial distribution of gene expression on UMAP plots for Ddx58 (RIG-I), Ifih1 (MDA5), and Cgas (cGAS) in DMSO– and pitavastatin-treated mice. Color intensity indicates the expression levels. H) Violin plots depicting the expression of selected genes across alveolar macrophage subclusters in DMSO (orange) and pitavastatin-treated (green) mice.

Trajectory analysis suggested a transition from cluster 0 to either cluster 7 or cluster 5, with cluster 0 potentially representing a progenitor or earlier differentiation state (Figure 6F). Furthermore, cluster 7 appeared to represent an intermediate state, subsequently resulting in further differentiation into clusters 1 and 2. It is notable that comparative UMAP and violin plot analyses showed that pitavastatin treatment increased the expression of RIG-I (DDX58) in clusters 3, 5, 8, and 10 even in the absence of poly I:C injection (Figures 6G and H). Moreover, MDA5 (IFIH1) expression was upregulated by pitavastatin treatment in clusters 8 and 10 (Figures 6G and H), and cGAS expression was enhanced in cluster 5 (Figures 6G and H). These single-cell analyses indicated that specific subsets of alveolar macrophages induced RIG-I expression in response to pitavastatin administration in vivo, and the cluster increasing the RIG-I expression was distinct from the cGAS expressing cluster.

To confirm the results obtained from the single-cell analyses, we first investigated whether total fraction of alveolar macrophages express RIG-I and noncanonical type I IFNs in response to pitavastatin administration in the absence of poly I:C stimulation. Alveolar macrophages were collected from pitavastatin-injected mice, and the expression of RIG-I and noncanonical type I IFNs was measured (Figure 7A). Since IFN-W1 gene is not conserved in mice, we examined the other noncanonical type I IFNs. As expected, the mRNA expression levels of RIG-I and noncanonical type I IFNs, including IFN-α5, –ε, and –ζ, were higher in alveolar macrophages isolated from pitavastatin-administered mice than those from control mice (Figure 7B). Subsequently, we sought to confirm the existence of subsets of alveolar macrophages corresponding to clusters 3, 6, 8, and 10 that can induce RIG-I expression in response to pitavastatin. Single-cell analyses determined the genes exclusively expressed in clusters 3, 6, 8, and 10 (Figure 7C). We performed flow cytometry to detect the alveolar macrophages expressing the cluster-specific markers. Interestingly, approximately 10% of alveolar macrophages expressed CD140a encoded by Pdgfra, which was exclusively expressed in cluster 6 (Figure 7D). Moreover, isolated CD140a^+^ alveolar macrophages markedly increased the expression of RIG-I in response to pitavastatin treatment in vitro (Figure 7E). These data provide evidence for the existence of a subset of alveolar macrophages whose RIG-I expression is enhanced by cholesterol restriction.

**Figure 7.**
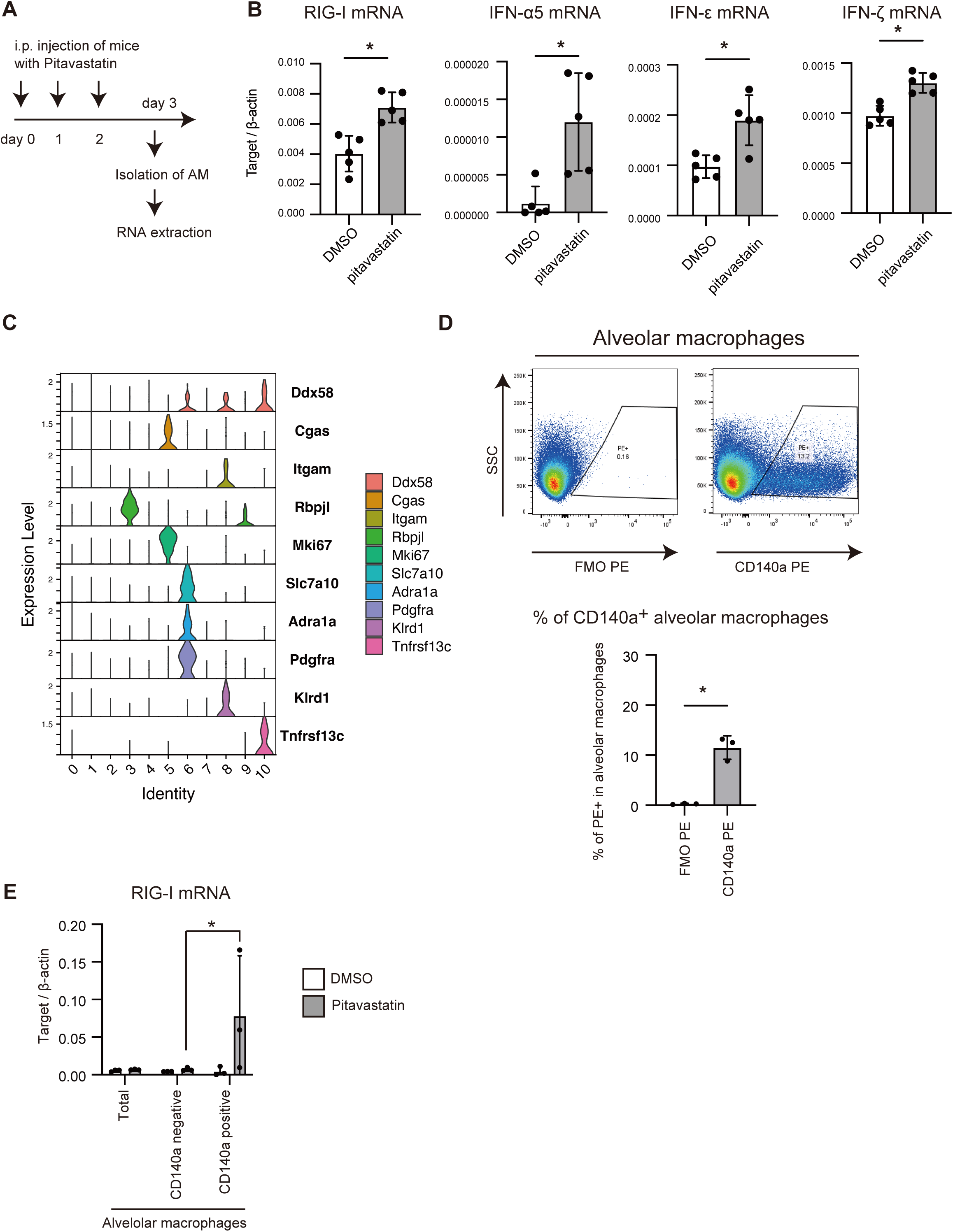
Priming of RIG-I pathway in CD140a^+^ alveolar macrophages. A, B) Mice were intraperitoneally injected with pitavastatin or solvent (DMSO) every three days, and alveolar macrophages were isolated (A). The expression of each gene in the isolated alveolar macrophages was determined by RT-qPCR and normalized to that of β-actin (B). C) A violin plot depicting cluster-specific gene expression in alveolar macrophages. D) Alveolar macrophages (CD45^+^, Siglect F^+^, and CD11c^+^) were isolated from the mouse bronchoalveolar lavage fluid (Figure EV7), and cells were analyzed by flow cytometry to observe cells expressing CD140a encoded by Pdgfra. E) CD140a positive alveolar macrophages were isolated from the lung using a MACS column, and cells were treated with DMSO or pitavastatin (2 μM) for 24 h. The expression of RIG-I mRNA was compared among CD140a positive, CD140a negative (flow-through), and total (CD140a positive + CD140a negative) alveolar macrophage fractions.

## Discussion

The link between cholesterol metabolism and the risk of disease severity and death due to viral infections has gained increasing attention, with mounting evidence supporting this correlation ^5, 29, 37^. Previous studies have illustrated the activation of the cGAS-STING pathway through the SREBP2 transcription factor following cholesterol depletion as well as the enhancement of RIG-I signaling by cholesterol limitation via MAVS protein geranylgeranylation ^3, 30, 31^. Extending upon these findings, our study elucidated another mechanism by which cholesterol limitation triggers a cell-type-specific induction of noncanonical type I IFNs, such as IFN-ω and IFN-α16. This process increased RIG-I expression, which, in turn, amplified RIG-I-dependent type I IFN expression. Our single-cell RNA transcriptome data corroborated that suppressing cholesterol synthesis increased the expression of RIG-I in specific subsets of alveolar macrophages in vivo. These findings indicate that restricting cholesterol synthesis primes the RIG-I pathway via noncanonical type I IFNs to enhance antiviral response in specific cell types. Considering the distinct expression of cGAS and RIG-I in the alveolar macrophage clusters, we anticipate that both cGAS and RIG-I are involved in the immunometabolism related to cholesterol synthesis.

The protein geranylgeranylation ligand GGPP, a metabolite of the mevalonate pathway that is inhibited by statin treatment, is associated with the innate immunity ^30, 38^. Since the geranylgeranylation of MAVS is inhibited by statins ^31^, this modification could be responsible for the statin-mediated enhancement of type I IFN expression. Indeed, a previous study reported that simvastatin enhanced IFN-β expression following Flu infection in macrophages ^31^. However, our study revealed that statin treatment can elevate the expression of noncanonical type I IFNs even in the absence of viral infection, a condition in which MAVS is not activated. Moreover, the expression of these noncanonical type I IFNs depended on the type I IFN receptor and SREBP1, and the absence of IFNAR2 or SREBP1 abolished the effect of statins in RIG-I-mediated IFN-β expression, as shown in Figures 3J and 4E. These observations indicate that the geranylgeranylation of the MAVS protein and the noncanonical type I IFN pathway cooperatively enhance RIG-I-dependent antiviral response in response to the restriction of cholesterol.

Viral infection induces the expression of canonical type I IFNs, such as IFN-β, via IRF3 and IRF7 transcription factors. Over ten type I IFN genes are encoded in the human and mouse genomes, and all type I IFNs activate the type I IFN receptor, comprising IFNAR1 and IFNAR2 ^39^. However, this present study revealed a noncanonical type I IFN expression pathway that depends on the SREBP1 transcription factor. This noncanonical type I IFN expression pathway does not require viral infection and can prime antiviral innate immune response, especially the RIG-I-dependent pathway, even in the absence of viral infection. In human cells, IFN-ω was regulated by the SREBP1. Unlike humans, the mouse genome does not encode IFN-ω. However, several noncanonical type I IFNs, such as IFN-ε, IFN-ζ, and IFN-α5, were upregulated in mouse alveolar macrophages in response to pitavastatin administration without viral infection. It remains unclear whether the SREBP1 is responsible for the mouse noncanonical type I IFN expression pathway; thus, further studies are required to elucidate the noncanonical pathways in different species.

Mouse alveolar macrophages can be identified using specific markers, such as CD11c, SiglecF, CD64, F4/80, and MerTK, and are conventionally classified into tissue-resident alveolar macrophages and monocyte-derived recruited alveolar macrophages ^40^. Recruited alveolar macrophages express relatively lower levels of Siglec F than tissue-resident alveolar macrophages ^40, 41^. Although our single-cell transcriptome analysis indicated that cluster 9 expressed lower levels of Siglec-F than the other clusters, it was unclear whether cluster 9 represented the recruited macrophages. Moreover, the single-cell transcriptome analyses revealed several clusters expressing RIG-I in response to pitavastatin. Flow cytometry revealed the presence of CD140a^+^ alveolar macrophages, and this macrophage fraction is expected to belong to cluster 6 and is distinct from cluster 5, which expresses cGAS and Mki67, which encodes the proliferative marker Ki67 ^42, 43^. The differential expression of RIG-I and cGAS implies that alveolar macrophages are heterogeneous cell populations comprising several functionally distinct populations. Thus, we expect that both the cGAS and RIG-I pathways synergistically augment antiviral innate immune responses, reflecting cholesterol levels in vivo.

Globally, we have repeatedly been confronted with pandemics caused by emerging viruses ^44^. Vaccines and medicines have been developed for various viral infectious diseases; however, these take time to develop to cope with new emerging viruses. Our findings indicate that cholesterol restriction or a limited amount of noncanonical type I IFNs primes RIG-I-dependent antiviral innate immune responses. Targeting this mechanism may be useful for establishing new treatment methods to cope with future viral pandemics.

## Methods

### Cell lines and cell culture

HeLa cells were cultured in Eagle’s MEM (Nissui) with 10% heat-inactivated fetal calf serum (FCS) and 10% L-Glutamine (Wako). HEK293 and HepG2 cells were cultured in low glucose DMEM (Wako) with 10% heat-inactivated FCS and penicillin/streptomycin (Wako). RAW264.7 and A549 cells were cultured in high glucose DMEM (Wako) with 10% heat-inactivated FCS and penicillin/streptomycin. THP-1 cells were cultured in RPMI (Wako) with 5% heat-inactivated FCS and penicillin/streptomycin. These cells were grown in a humidified incubator maintained at 37 ℃ with a 5% CO₂ atmosphere. IFNAR2, SREBP1, and SREBP2 KO cells were generated using the CRISPR/Cas9 system. The guide sequence was cloned into the BbsI restriction site of the pX459 plasmid, which encodes Cas9 and puromycin-resistance genes. The plasmid carrying the guide sequence was then transfected into HeLa cells. After 36 h, the cells were treated with 1 μg/mL of puromycin for 3 days to isolate the cells with the plasmids. Then, cells were then cultured in fresh medium without puromycin for 2 days and were seeded onto 96-well plates to obtain single clones. The guide sequence and purchased reagents are listed in Table S1. Pitavastatin and cerivastatin were dissolved in DMSO at a concentration of 10 mM. Cells were treated with DMSO concentrations of 0.1 % or less. The treatment of cells with pitavastatin was performed at 2 μM for 24 h, unless otherwise stated.

### Screening of compounds

A validated compound library was provided by the Drug Discovery Initiative (The University of Tokyo). In the first screening, HEK293 cells were transfected with pEF-BOS-AcGFP, p125-mCherry, pEF-BOS-FLAG-tagged-RIG-I, and pEF-BOS-HA-tagged-Riplet plasmids using Lipofectamine 2000, treated with each compound simultaneously, and cultured in 96-well plates. After 48 h, the fluorescence of AcGFP and mCherry was observed using a BZ-X700 microscope (Keyence), and fluorescence intensities were measured with BZ-H1A software (Keyence). The promoter activity of the IFN-β gene was calculated by dividing the fluorescence levels of mCherry by those of AcGFP. In the second screening, HEK293 cells were transfected with p125-Luciferase, pRL-TK-Renilla, pEF-BOS-FLAG-tagged-RIG-I, and pEF-BOS-HA-tagged-Riplet expression plasmids using Lipofectamine 2000 and treated with each compound, which were isolated by the first screening, and subsequently cultured in white 96-well plates (Corning). Cell lysates were prepared 24 h after transfection, and the activities of luciferase and *Renilla* luciferase were determined using the Dual-Luciferase Reporter Assay system (Promega). The luminescence was measured using a FilterMAX F5 microplate reader (MOLECULAR DEVICES), and the promoter activity of IFN-β was determined by normalizing luciferase activities to *Renilla* luciferase activities.

### Mice

C57BL/6JJcl mice were purchased from Kyudo. Leptin KO (ob/ob) mice and control WT mice were purchased from Jackson Laboratories, and maintained at the Kumamoto University Center for Animal Resources and Development (CARD) in a specific pathogen-free (SPF) environment. All experimental procedures were approved by the Institutional Animal Committee of Kumamoto University and performed according to their guidelines. To isolate the alveolar macrophages, the lungs were flushed with 800 μL BAL buffer (PBS with 2 mM EDTA and 0.5% EDTA) at least five times. The collected BAL fluid (BALF) was centrifuged at 400 g.

### Cholesterol concentration

Cellular and mouse serum cholesterol levels were measured using the Cholesterol/Cholesterol Ester-Glo Assay Kit (Promega). Cells were cultured in 96-well plates and then treated with the compounds at various concentrations. The cells were gently washed with PBS three times, and cell lysates were prepared according to the manufacturer’s protocol. Blood was collected from the tail vein of each mouse and allowed to stand for 30 minutes in tubes. Incubated blood was centrifuged at 1,300g, and then serum was collected. The serum was diluted 50–80 times using the reagents provided in the kit. Diluted samples were dispensed into 96-well plates, and then experiments were performed according to the manufacturer’s instructions. Luminescence level was measured using a FilterMAX F5 microplate reader, and cholesterol concentrations were determined based on standard curves.

### Microarray analysis

HeLa cells were cultured in 24-well plates and treated with DMSO or pitavastatin. After 24 h, the cells were infected with SeV at a multiplicity of infection (MOI) of 10 for 24 h. Total RNA was isolated using TRIZOL reagent. cDNA and Amino Allyl aRNA were synthesized using the Amino Allyl MessageAmp II aRNA Amplification Kit (Ambion#1753). CyeDye coupling and fragmentation were performed according to the protocol supplied by TORAY. Hybridization was performed for 16 hr at 37°C using a rotary shaker (set at 250 rpm). The hybridization buffer and washing protocol were performed according to the protocol provided by TORAY. A 3D-Gene scanner (TORAY) was used for scanning. Images were quantified by extraction (TORAY). The raw data for each spot were normalized by substituting it with the mean intensity of the background signal, which was determined using the signaling intensities of all blank spots within 95% confidence intervals. The microarray data were deposited in GEO under the accession number GSE199783.

### Western blotting

Cells were washed with PBS and lysed with lysis buffer (20 mM Tris-HCl pH7.5, 125 mM NaCl, 1 mM EDTA, 10% glycerol, 1% NP-40, 30 mM NaF, 5 mM Na_3_VO_4_) in the presence of a protease inhibitor cocktail (Sigma Aldrich). Cell lysates were placed on ice for 30 min and were subsequently centrifuged for 20 min at 15,000 rpm at 4°C. The supernatants were transferred to 1.5 mL tubes. Protein levels were measured using the BCA Protein Assay Kit (TaKaRa). β-mercaptoethanol was added to cell lysates, which were subsequently boiled for 5 min at 95 ℃. Samples and protein markers (Bio-Rad) underwent Tris-glycine SDS-polyacrylamide gel electrophoresis (SDS-PAGE) and were transferred onto PVDF membranes. The membrane was blocked with 5% skim milk in rinse buffer (0.1% tween 20, 10 mM Tris-HCl at pH7.5, 0.8% NaCl, and 1mM EDTA) and subsequently incubated with primary antibody (1:1000) at 4 °C overnight, and then incubated with HRP-conjugated secondary antibody (1:10000) for 60 min at room temperature. Immunoblots were visualized using Amersham ECL prime western blotting detection reagent (GE Healthcare) and detected with a Bio-Rad ChemiDoc Touch imaging system. The antibodies used for western blotting are listed in Table S1.

### Reporter assay

HEK293 cells were cultured in 24-well plates and transfected with the indicated reporter and protein expression plasmids together with the pEF-BOS-Renilla plasmid by using Lipofectamine 2000. Cell lysates were prepared 24 h after transfection, and luciferase and Renilla luciferase activities were determined using the Dual-Luciferase Reporter Assay system (Promega). The luminescence level was measured using a luminometer (ATTO), and the promoter activity was determined by normalizing luciferase activity to *Renilla* luciferase activities.

### ELISA

Cells were cultured in 24-well plates and treated with pitavastatin for 24 h, and infected with SeV or Flu for 24 h. Cell culture supernatants were collected and analyzed for IFN-β using an ELISA kit (Verikine Human IFN-Beta ELISA kit) (PBL). Serum was collected from the mouse tail vein and IFN-β protein levels were measured with the ELISA kit according to the manufacturer’s instructions.

### RT-qPCR

Total RNA was isolated using TRI reagent (MOR), according to the manufacturer’s protocol and then treated with DNase I. cDNA was generated using a high-capacity cDNA Reverse Transcription Kit (Life Technologies). RT-qPCR was performed with Luna Universal qPCR Master Mix (Biolabs) using the Step-One Real-time PCR system (Life Technologies). Target mRNA expression was normalized to the expression of GAPDH or β-actin, as indicated. The fold increase was calculated by dividing the mRNA levels at the indicated time points by those at 0 h or by dividing the mRNA levels following pitavastatin treatment with the vehicle. The primer sequences are listed in Table S1.

### ChIP-qPCR

4 × 10^6^ of HEK293 cells were cultured in a 10 cm dish and transfected with the indicated FLAG-tagged protein expression vectors. After 24 h, cells were subjected to a ChIP assay, according to the manufacturer’s instructions (Cell Signaling Technology number 9003). The cells were sonicated 15 times, at 30 sec intervals. Immunoprecipitation was performed using an anti-FLAG antibody and control IgG. The qRT-PCR was performed using the primers listed in Table S1.

### 10X Single-cell RNA sequencing

Pitavastatin calcium was dissolved in DMSO to achieve a concentration of 25 mg/mL. 13-week-old female mice were intraperitoneally injected with 8 μL of DMSO or pitavastatin calcium solution in 200 uL of PBS (final pitavastatin dosage: 10 mg/kg/day) for three days. The lung tissues were harvested, snap-frozen in liquid nitrogen, and stored at –80 ℃. The lung tissue was fixed with formaldehyde and dissociated using a GentleMACS Octo Dissociator (Miltenyi Biotec). After hybridizing the detection probe from the chromium fixed RNA kit for the mouse transcriptome (10x Genomix) with intracellular RNA, a ligation reaction was performed to join the hybridized probes. After mixing each sample to ensure the number of cells was equal, the barcoded primer beads and cells were encapsulated using the Chromium Next GEM Chip Q Single Cell Kit on the Chromium X platform (10x Genomix). The probe and primer were annealed, and an extension reaction was performed to create a detection probe. Thereafter, a sequence library was prepared by PCR amplification and purification using the indexed primers included in the Dual Index Kit TS Set A (10x Genomix). A quality check of the sequence library was performed with an Agilent 2100 BioAnalyzer using a high-sensitivity DNA kit. Libraries were sequenced on an Illumina NovaSeq X Plus in pair-end mode. Library preparation and sequencing after tissue fixation were performed by Takara Bio Inc.

### Processing of the scRNA-seq data set

Raw sequencing data were processed and the reads were aligned to the mouse genome (mouse mm10; refdata-gex-mm10-2020-A). Following this, gene expression was quantified using Cell Ranger (v.7.1.0) software. The expression matrices were processed in R (v.4.3.1) using Seurat (v.5.0.1) as described below. We only selected the genes whose expression (unique molecular identifiers; UMI ≥ 1) was detected in >3 cells. Cells with more than 200 detected UMIs were retained, and filtering was applied (≥ 200 or < 5,000 genes per cell and % mitochondrial UMI counts < 5). In total, 41,259 cells (DMSO group: 18,415 cells and pitavastatin group: 22844 cells) were selected for further analysis out of a total of 42,606 cells (DMSO group: 19,024 cells and pitavastatin group: 23,582 cells). Data were normalized, scaled, and subjected to principal component analysis (PCA) using 2,000 highly variable genes identified using the Seurat vst algorithm. We used the IntegrateLayers function with the HarmonyIntegration method to integrate different samples without the batch effect. The RunUMAP, FindNeighbors, and FindClusters functions were applied for clustering, with 30 dimensions identified using Harmony at a 0.5 resolution. For automatic cell type annotation, we used Single R (v2.2.0) software and LungMAP_MouseLung_CellRef (celltype_level3) ^45^ data sets. For extracting the alveolar macrophage subcluster, we chose cells defined as “AM” by LungMAP_MouseLung_CellRef and applied another filter using CellSelector software. The marker genes specifically expressed in each cluster were identified using the FindAllMarkers function and represented using a dotplot and heatmap using the Seurat function. To perform cell cycle scoring, a list of cell cycle markers was loaded, and the CellCycleScoring function in Seurat was employed. The Monocle 3 software was used for trajectory analysis of the alveolar macrophage. To analyze the mean expression level of given gene signatures, we applied AddModuleScore functions in addition to a calculation method described in previous reports ^46, 47^. Using the abovementioned methods, the normalized and log-scaled expression value of each gene was min-max scaled to adjust each gene to have an equal dynamic range (max = 1.0, min = 0). The mean expression levels of the gene set for each single cell were then calculated.

## Statistical analysis

Error bars represent the standard deviation (SD). Statistical significance (p-value) was determined using a two-tailed Student’s *t*-test, one-way ANOVA, or two-way ANOVA using Prism ver. 10 (GraphPad Software)

## Data availability

Microarray and scRNA-seq data were deposited into the GEO database, and the accession numbers were GSE199783 and GSE264004, respectively.

## Acknowledgments

We thank our laboratory members for their helpful discussion and technical assistance. This work was supported in part by the Japan Society for the Promotion of Science (JSPS, Grant No.: 23H02949), the Japan Agency for Medical Research and Development (AMED, Grant No. 23gm1610011h), the Kose Cosmetology Foundation, Takeda Science Foundation, and the Uehara Memorial Foundation.

## Author contributions

TN conducted the experiments. AO examined the effect of cerivastatin. TN prepared the samples for single-cell transcriptome analysis and the obtained data was analyzed by KT and YM. TK and HO designed the experiments. TN, TK, KT, and HO wrote the manuscript.

## Expand View Figure legends

**Figure EV1. Effects of statins on HeLa cells**

A–G) HeLa cells were treated with pitavastatin or DMSO (mock) for 24 h, Subsequently, they were stimulated by either transfecting short poly I:C (A, B), infection with SeV (C), adding poly I:C to cell culture medium (D, E), transfecting cGAMP (F), or being infected with Flu (G). The expression of each gene was determined by RT-qPCR and normalized to that of GAPDH.

H–J) HepG2 (H), A549 (I), and HeLa (J) cells were treated with pitavastatin (H, I), cerivastatin (J), or solvent (DMSO) for 24 h, and the cellular cholesterol levels were measured. Statin-pretreated cells were infected with SeV for 24 h, and then the expression of IFN-β mRNA was determined by RT-qPCR and normalized to that of GAPDH.

K, L) HeLa (K) and HepG2 (L) cells were transfected with HMGCR or control siRNA. 24 h after siRNA transfection, the cells were stimulated by short poly I:C transfection for the indicated time period. The mRNA expression of each gene was determined using RT-qPCR and normalized to that of GAPDH.

**Figure EV2. The expression of non-canonical type I IFNs after pitavastatin treatment**

A) HeLa cells were treated with pitavastatin for 24 hr. The expression of each gene was determined by RT-qPCR and normalized to that of β-actin. The fold increase was calculated by dividing the normalized expression level of a pitavastatin-treated sample by a pitavastatin-untreated sample value.

B) HEK293, HepG2, A549, THP-1, and HeLa cells were treated with pitavastatin for 24 h, and IFN-ω expression was determined by RT-qPCR and normalized to that of GAPDH.

**Figure EV3. Effect of IFN-ω on the RIG-I-dependent innate immune response**

A) HeLa cells were treated with 100 pg/mL of recombinant IFN-ω protein for 24 h, and the expression of each gene was determined by RT-qPCR and normalized to that of GAPDH.

**Figure EV4. The role of SREBPs in type I IFN response**

A) HEK293 cells were transfected with a p125-luc (IFN-β promoter) or IFN-ω promoter reporter plasmid along with a *Renilla* luciferase-expressing vector (internal control) and MAVS, MyD88, STING, or SREBP1a-N expressing vector. 24 h after transfection luciferase activities were measured and normalized to *Renilla* luciferase activity.

B–E) HEK293 cells were transfected with 200 ng/mL of RIG-I CARD expressing vectors or 200, 1000, or 2000 ng/mL of SREBP1a-N, 1c-N, or 2a-N expressing vectors in a 24-well plate. The total amount of DNA was adjusted to the same amount by adding an empty vector. 24 h after transfection, the expression of each gene was determined by RT-qPCR and normalized to that of β-actin.

F) HeLa cells were stimulated via short poly I:C transfection, and the mRNA expression of each gene was determined by RT-qPCR. The fold increase was calculated by dividing mRNA expression at each time point by that at 0 h.

G) THP1 cells were stimulated via transfection with short poly I:C, and the expression of IFN-ω mRNA was determined by RT-qPCR and normalized to that of GAPDH.

H, I) Promoter regions of IFN-ω and IFN-α16. The E-box and SRE motifs are shown in the green and yellow boxes, respectively. Numbers represent the positions of the nucleotides from the start of the ORF.

**Figure EV5. The tissue-specific effect of pitavastatin on type I IFN responses**

A) Pitavastatin or solvent (DMSO) was intraperitoneally injected into mice every day for three days, followed by intraperitoneal injection of poly I:C or solvent (DMSO). 24 h after poly IC injection, the lung, heart, muscle, and uterus were isolated. The total RNAs was extracted from each tissue. The expression of each gene was determined by RT-qPCR and normalized to that of β-actin.

**Figure EV6. Single-cell RNA-seq analysis of mouse lungs following pitavastatin administration**

A) Schematic representation of the single-cell analysis. Mice were injected with pitavastatin or solvent (DMSO) every three days. The lung tissues were harvested and subjected to single-cell RNA-Seq analysis.

B) UMAP visualization of total lung cell populations identified in the scRNA-seq data. Each dot represents a single cell, with colors indicating different cell types described in the legend to the right of the figure.

C) UMAP plots comparing the distribution of lung cell populations between pitavastatin and DMSO-injected mice. The bottom bar graph shows the percentage of each cell type in the respective groups.

D) Functional annotation of gene expression profiles in lung cells from DMSO and pitavastatin-injected mice. The top UMAP plots highlight cells with expressing genes related to the cholesterol metabolic process (GO: 0008203), while the bottom plots show cells with expressing genes related to type I IFN production (GO: 0032606). The intensity of color intensity corresponds to the expression level of the gene associated with the respective biological process.

**Figure EV7. Gating strategy for the isolation of alveolar macrophages**

Cells were isolated from wild-type mice lungs. The gating strategies for the identification of alveolar macrophages were shown.

